# Translocation and duplication from CRISPR-Cas9 editing in *Arabidopsis thaliana*

**DOI:** 10.1101/400507

**Authors:** Peter G. Lynagh, Soichi Inagaki, Kirk R. Amundson, Mohan P.A. Marimuthu, Brett Randolph Pike, Isabelle M. Henry, Ek Han Tan, Luca Comai

## Abstract

Cut DNA ends in plants may recombine to form novel molecules. We asked whether CRISPR-Cas9 expression in plants could induce nonhomologous recombination between diverse and heterologous broken DNA ends. We induced two breaks separated by 2.3 or by 8.5 kilobases leading to duplication of the intervening DNA and meiotic transmission of the 2.3kb duplication. Two or more dsDNA breaks in nonhomologous chromosomes led to ligation of breakpoints consistent with chromosome arm translocations. Screening 881 primary transformants we obtained 195 PCR products spanning independent, expected translocation junctions involving ends produced by cutting different loci. Sequencing indicated a true positive rate of 84/91 and demonstrated the occurrence of different junction alleles. A majority of the resulting structures would be deleterious and none were transmitted meiotically. Ligation of interchromosomal, heterologous dsDNA ends suggest that the CRISPR-Cas9 can be used to engineer plant genes and chromosomes in vivo.

**Significance Statement:** We explored how genome editing tools such as CRISPR-Cas9 could provide new ways to tailor novel genomic combinations and arrangements. We show that distant cut ends often precisely come together, that cuts in different chromosomes can result in translocations, and that two cuts within a chromosome often result in the duplication of the intervening segment. Formation of multiple structures with precise junctions will enable engineered rearrangements that can be predicted with accuracy.

## Introduction

The discovery of CRISPR-Cas9 (for Clustered Regularly Interspaced Short Palindromic Repeats and CRISPR-associated protein 9) and related systems is enabling precise editing of plant genomes ^1^. With single base pair precision, CRISPR-Cas9 is capable of making a blunt double stranded break (DSB) in the DNA of living cells ^2,3^. The Cas9-sgRNA (single-guide RNA) complex scans the genome for Protospacer Adjacent Motif (PAM) sequences, which is predominantly the sequence NGG for Cas9 ^4^. Once bound to the PAM sequence, upstream base pairs are sequentially opened by the complex and tested for complementarity to a programmable 20 bases within the sgRNA ^5^.

Imprecise repair of DSB within genes results in small indels and loss of gene function, as demonstrated in *Arabidopsis thaliana* ^6,7^. Efficient expression of the sgRNA and Cas9 constructs should increase editing probability. In most studies, *Arabidopsis* Pol-III snRNP U6-26 promoter seems adequate for sgRNA expression ^6–10^. Multiple gRNA can be expressed by driving each one with its own U6-26 ^11^, or by post-transcriptional processing of a gRNA polycistron driven by U6-26 ^12,13^ or a Pol-II promoter such as that from Cestrum yellow leaf curling virus ^11,14^.

Different expression strategies have been used for Cas9 with the goal of obtaining a higher percentage of germline mutations. Many changing variables and approaches make it difficult to compare results, and mutation efficiencies have varied greatly. This may be explained by the observation that Cas9 endonucleolytic activity is directly related to the concentration of the ribonucleoprotein ^15^. Driving Cas9 expression with CaMV 35S promoter, which has been effective in other transgenic applications ^16^, produced a surprisingly low mutation efficiency in the germline and often also in somatic tissues ^7,8,10,17,18^. This has stimulated the testing of promoters active in reproductive and meristematic cells, such as those driving *EC1.2*, *UBQ1*, *UBQ10*, *YAO*, *INCURVATA2*, *SPOROCYTELESS* and *RPS5A* ^7–10,18–20^. In some cases, germline mutation frequencies are lower than somatic mutagenesis frequencies, such as with *YAO* and *INCURVATA2* promoters ^19,20^. Nonetheless, several of these promoters proved superior to CaMV 35S for inducing mutations. For example, Tsutsui and Higashiyama found a 2nd leaf albino phenotype in 3/96 (3%) Arabidopsis primary transformants when using the CaMV 35S promoter and 38/57 (67%) when using the RPS5A promoter. This improved efficiency should help to expand the potential of the CRISPR-Cas9 system to produce heritable knock-out mutations.

In addition to gene knock-outs, DSBs could lead to new DNA structural arrangements useful in basic studies and for plant chromosome engineering, but unwanted if the objective is a simple knockout ^21^. Each Cas9-induced DSB produces two free ends that become actively managed by the plant’s own elaborate DSB repair processes. The plant typically reconnects the two cut ends through the Nonhomologous End Joining (NHEJ) DSB repair pathway ^1,22,23^. The cut and repair process may go through many cycles until the religated ends are mutated, preventing recognition and cutting by the Cas9-sgRNA complex ^14^. When two DSBs are made close to each other on a single chromosome, segment deletions are frequent and combinatorial outcomes have been exploited to generate quantitative alleles ^24^. Deletions are often accompanied with perfect ligation of the two flanking ends. In the case of cuts spaced 40-80bp, inversions of the intervening fragment have been also observed ^14^. If distant DSBs are made on a single chromosome, intrachromosomal recombination may occur, as has been demonstrated by CRISPR-induced deletions fifty to hundreds of kb long ^14,25^. Intrachromosomal recombination is likely facilitated by the existence of chromosome-specific nuclear domains, discovered in 3-D conformation analysis of chromatin ^26^. Nonetheless, in animals, interchromosomal translocations have been documented as well ^27–30^. Targeted duplications of various sizes have also been produced ^31^. The duplications were proposed to be produced by allelic recombination or segmental translocation.

Not all possible recombination products have been reported in plants. *Trans* cuts produced by the rare-cutting endonuclease *I-SceI* produced chromosomal translocation ^32^. In addition, low frequency and imprecise translocations were observed after CRISPR-Cas9 *trans* cuts ^33^. Duplications that are dozens of base pairs long have been produced ^34^, but it would be interesting if gene-sized segments could be duplicated. In addition to finding interchromosomal translocations, a tandem duplication was briefly mentioned as a result from treating tetraploid *A. thaliana* with a restriction enzyme that is estimated to cut every 256 bp <u>(Muramoto et al. 2018)</u>. A cut segment may also self-ligate, forming a circle, a structure that could potentially amplify if endowed with replication signals.

In order to make optimal use of this tool, it is important to document what type of intra-and interchromosomal cut ends can efficiently find each other before religation and mutagenesis stops the CRISPR endonucleolytic process. Pursuing this objective, we confirmed high germline mutation efficiency when using the RPS5A promoter to drive Cas9 activity ^10^. Indeed, on some targets we observed >85% efficiency in the soma and germline. Using material where cutting is demonstrably frequent, we analyzed the structural range of mutations that result from the use of proRPS5A-Cas9 (RC9) in *Arabidopsis thaliana*. We show that after cutting multiple targets, disparate heterologous ends can join in perfect unions forming predictable duplications and translocations.

## Results

### The RPS5A promoter is active in the zygote and early embryo

We reasoned that the induction of dsDNA cleavage as early as possible during embryo development would enhance germinal transmission, avoid endopolyploidy, increase the frequency of whole-plant modifications, while taking advantage of DNA repair pathways that are most active in meristems ^35–37^. Therefore, we chose the promoter of the gene encoding the 40S ribosomal protein S5 (*RPS5A*, gene *AT3G11940*) because of its activity in the early embryo and proliferating cells ^38^. To confirm the previously published expression pattern, we used the *RPS5A* promoter to drive expression of the td-Tomato fluorescent protein ^39^ fused to histone protein H2B in *A. thaliana.* Expression was observed in the zygote (n=42), two-cell (n=30) and four-cell embryos (n=31) (Fig. 1) as well as at later stages (n=8, data not shown). Consistent with previous reports ^10,38^, the fluorescent nuclei indicate that transgenes driven by the RPS5A promoters are active both zygotically and in later proliferative stages.

**Figure 1.**
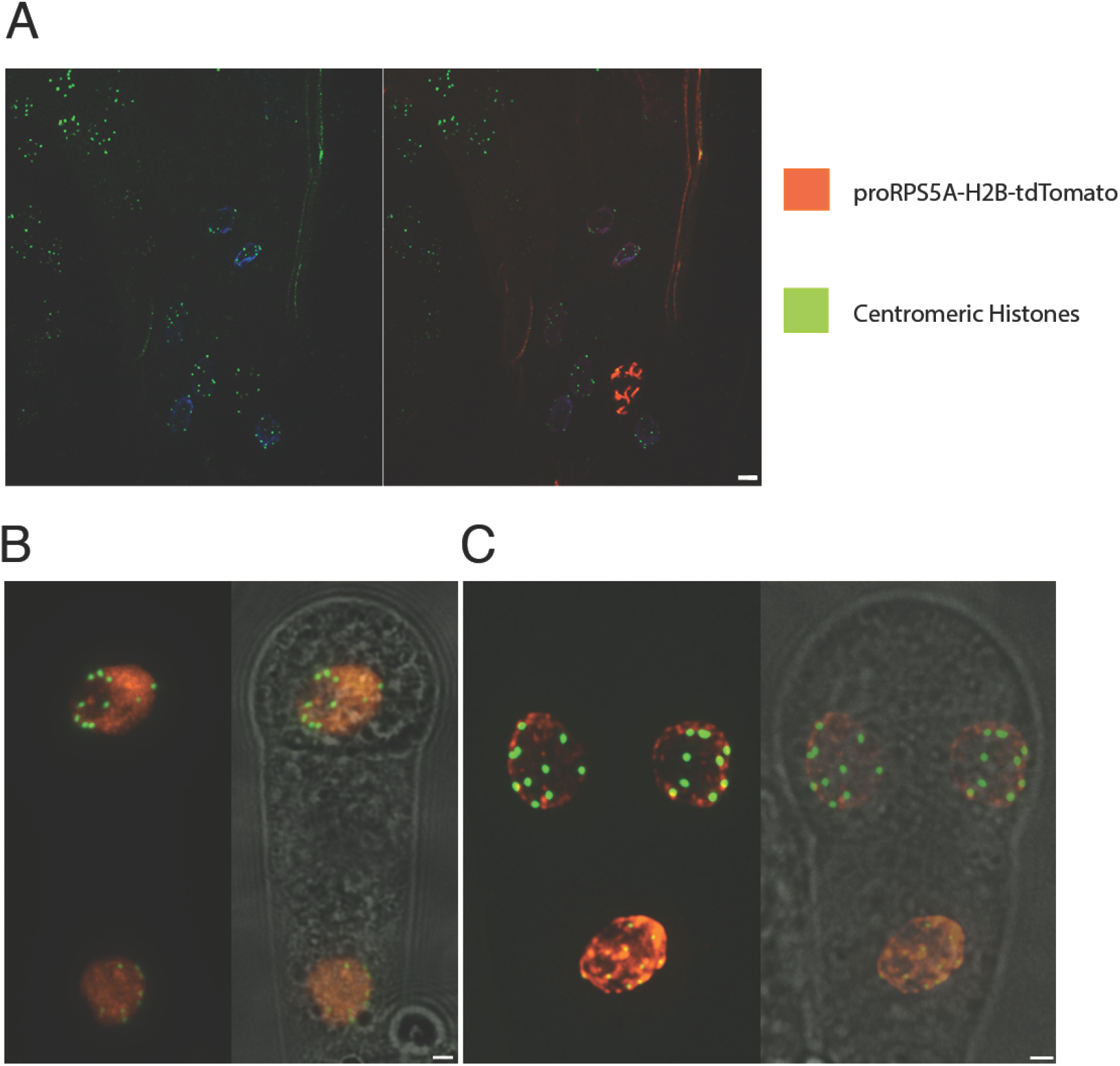
*RPS5A-Cas9* drives efficient expression in zygotic and early embryonic cells. The *RPS5A* promoter expresses a red fluorescent td-Tomato fused to histone 2B in the *Arabidopsis* zygote, two-cell embryo and three-cell embryo. A plant expressing *proCENH3-GFP-CENH3[TAILSWAP]* was pollinated by a plant expressing *RPS5A-td-Tomato-H2B*. A. Egg sac after fertilization showing condensing red-stained chromatin. B. Two and (C) three-cell embryos. Red: nuclear chromatin containing *td-Tomato-H2B*. Green: the punctate signal corresponds to centromeres stained by deposition of centromeric fusion protein *GFP-CENH3[TAILSWAP]*. Blue: DAPI DNA stain. Scale = 10μm

### Efficient editing of genes by RPS5A promoter driven Cas9 (RC9)

Next, we assessed efficiency and developmental timing of CRISPR-induced mutations when Cas9 is driven by the *RPS5A* promoter, a cassette we named RC9. We first targeted *BRASSINOSTEROID-INSENSITIVE 1* (*BRI1*; *At4G39400*), whose homozygous knockouts exhibit a dwarf phenotype ^40^. Separately, we transformed *Arabidopsis* Col-0 with the RC9 transgene and with a previously published sgRNA ^7^ expressed under proAtU6-26. By providing Cas9 from one parent and the guide RNA from the other parent (see Methods), early CRISPR activity was evident. In the best parental combinations, we observed the widespread presence of dwarfs (41.2%; n=98) among hybrid seedlings (Table 1. Table S8). We concluded that this CRISPR-Cas9 system resulted in biallelic mutation of the wild-type targets in very early embryos or has widespread activity in most cells of the body.

**Table 1:** Summary of translocation characteristics and frequencies in T1 plants that each receivd 2 sgRNAs targeting estimated boundaries of a promoter and 1 sgRNA targeting a different chromosome. 1 Each PCR primer pair designates a translocation junction of interest. The primer locations are as described in Figure 7A. 2 Some targets were chosen so that a perfect translocation junction will reform a target sequence that is expected to be recut until an imperfect translocation junction is formed. (See Fig. S6). 3 Some targets were chosen so that a perfect circularization junction will reform a target sequence that is expected to be recut until an imperfect circularization junction is formed (See Fig. S6) 4 Segmental translocations are expected to result in monocentric translocations, but chromosome arm translocations can result in acentric, monocentric or dicentric chromosomes 5 Every locus in the Arabidopsis genome has a closest distance to the KNOT as measured in base pairs 48–50. The distance of each of the two target loci to the KNOT were added together as an estimation of the physical closeness of the two loci 6 Only junction CF is expected to activate *LEC1* or *WUS1*, because only that junction has the 3’ end of the highly expressed promoter facing into the 5’ end of *LEC1* or *WUS1* near their transcription start sites (TSS). Primer BE is noninformative regarding gene activation because that junction involves neither the 3’ promoter end nor the TSS of interest.

| Targets | PCR<sup>1</sup><br>Primer<br>Pair | Perfect<sup>2</sup><br>Junction<br>Recut? | Perfect<sup>3</sup><br>Circle<br>Recut? | Expected<sup>4</sup><br>Centromeres | Distance<sup>5</sup><br>through<br>KNOT<br>(kb) | Gene<sup>6</sup><br>Activation<br>Junction? | PCR<br>positive<br>T1s | # T1s<br>Tested | Freq. |
| --- | --- | --- | --- | --- | --- | --- | --- | --- | --- |
| Rabe1b-1<br>Rabe1b-2<br>Lec1-3 | BE | No | Yes | 1 | 700 | - | 4 | 70 | 0.06 |
|  | BF | No | Yes | 0 | 700 | No | 2 | 61 | 0.03 |
|  | CE | No | Yes | 2 | 700 | No | 0 | 70 | 0.00 |
|  | CF | No | Yes | 1 | 700 | Yes | 0 | 70 | 0.00 |
| Rabe1b-1<br>Rabe1b-3<br>Lec1-3 | BE | No | No | 1 | 700 | - | 0 | 46 | 0.00 |
|  | BF | No | No | 0 | 700 | No | 0 | 41 | 0.00 |
|  | CE | No | No | 2 | 700 | No | 0 | 60 | 0.00 |
|  | CF | No | No | 1 | 700 | Yes | 3 | 44 | 0.07 |
| Rabe1b-1<br>Rabe1b-3<br>Wus1-1 | BE | No | No | 0 | 4300 | - | 1 | 12 | 0.08 |
|  | BF | No | No | 1 | 4300 | No | 1 | 12 | 0.08 |
|  | CE | No | No | 1 | 4300 | No | 2 | 12 | 0.17 |
|  | CF | No | No | 2 | 4300 | Yes | 1 | 12 | 0.08 |
| PDE334-1<br>PDE334-4<br>LEC1-3 | BE | Yes | Yes | 2 | 700 | - | 1 | 16 | 0.06 |
|  | BF | No | Yes | 1 | 700 | No | 3 | 62 | 0.05 |
|  | CE | No | Yes | 1 | 700 | No | 7 | 101 | 0.07 |
|  | CF | Yes | Yes | 0 | 700 | Yes | 25 | 163 | 0.15 |
| PDE334-3<br>PDE334-2<br>LEC1-3 | CE | No | Yes | 1 | 700 | No | 0 | 41 | 0.00 |
|  | CF | No | Yes | 0 | 700 | Yes | 37 | 109 | 0.34 |
| PDE334-1<br>PDE334-2<br>LEC1-3 | BE | No | No | 2 | 700 | - | 2 | 121 | 0.02 |
|  | BF | No | No | 1 | 700 | No | 0 | 30 | 0.00 |
|  | CE | Yes | No | 1 | 700 | No | 0 | 72 | 0.00 |
|  | CF | No | No | 0 | 700 | Yes | 36 | 121 | 0.30 |
| PDE334-1<br>PDE334-4<br>WUS1-1 | BF | No | Yes | 2 | 4300 | No | 2 | 74 | 0.03 |
|  | CE | No | Yes | 0 | 4300 | No | 22 | 74 | 0.30 |
|  | CF | Yes | Yes | 1 | 4300 | Yes | 4 | 74 | 0.05 |
| PDE334-3<br>PDE334-4<br>WUS1-1 | BE | Yes | No | 1 | 4300 | - | 2 | 12 | 0.17 |
|  | BF | No | No | 2 | 4300 | No | 4 | 30 | 0.13 |
|  | CE | No | No | 0 | 4300 | No | 2 | 30 | 0.07 |
|  | CF | No | No | 1 | 4300 | Yes | 4 | 30 | 0.13 |
| PDE334-3<br>PDE334-2<br>WUS1-1 | BE | No | Yes | 1 | 4300 | - | 5 | 30 | 0.17 |
|  | BF | No | Yes | 2 | 4300 | No | 5 | 30 | 0.17 |
|  | CE | No | Yes | 0 | 4300 | No | 4 | 44 | 0.09 |
|  | CF | No | Yes | 1 | 4300 | Yes | 1 | 14 | 0.07 |
| PDE334-1<br>PDE334-2<br>WUS1-1 | BE | No | No | 1 | 4300 | - | 0 | 39 | 0.00 |
|  | BF | No | No | 2 | 4300 | No | 2 | 39 | 0.05 |
|  | CE | Yes | No | 0 | 4300 | No | 9 | 39 | 0.23 |
|  | CF | No | No | 1 | 4300 | Yes | 0 | 32 | 0.00 |

The *bri1* dwarfs are tedious to grow and to analyze further. In order to better understand heritability of the mutant phenotype, we next targeted the *CHLORINA1* gene *(CH1*; *At1G44446)*. *CH1* homozygous knockouts give rise to yellow-green plants. In this case, we targeted *CH1* with two different guide RNAs simultaneously and expressed CAS9 and sgRNAs from the same vector. One sgRNA targeted five base pairs upstream of the CH1 start codon (site *I*) and the other sgRNA targeted a conserved domain (PFAM PF08417) encoded by the ninth exon (Site *II*, Fig. 2-A). Deletion of the intervening fragment and frameshifting indels at site *II* should knock out function. The overall effectiveness of mutagenesis was demonstrated by the fact that 37.1% (13/35) of the T1 (primary transformant) plants were yellow and 9% (3/35) were half yellow and half green (Fig. 2-B). We phenotyped the selfed progeny of seven yellow T1 plants. In two families we found green individuals. Analysis of one of these two families, described below, indicates that biallelic knockout observed throughout the T1 body was not early enough to affect the germline.

**Figure 2.**
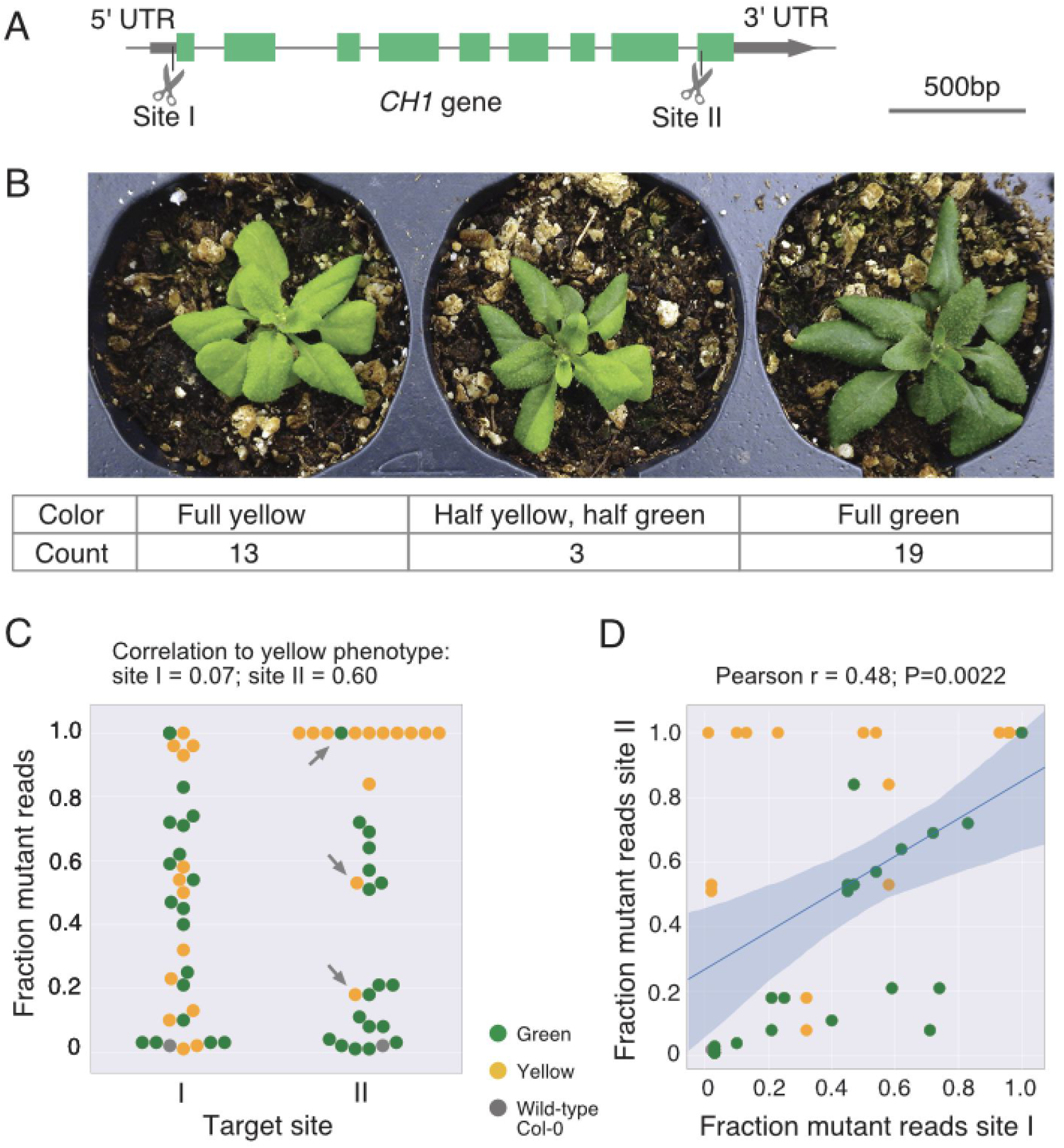
Analysis of *ch1* phenotypes induced by CRISPR-Cas9 in first generation transgenic plants. A. Structure of *CH1* gene and targeted sites. B. Typical phenotypes in a population of T1 individuals (primary transformants, n=39) with corresponding counts. C. Swarmplot summarizes the results of the phenotypes and Ampliseq analysis. Each dot represents a single T1 individual, with its phenotype illustrated by color. The Y-axis represents the fraction of AmpliSeq reads that were mutant. On the the X-axis are the two targeted sites in *CH1.* Mutations at site I affect non-coding exon DNA, are expected to be mostly neutral and uncorrelated to phenotype. Mutations at site II affect the conserved coding region, are deleterious and well correlated with phenotype. Arrows point at the major exceptions: a single green plant (#16, Table S4) with 100% mutated *CH1* site II. Upon further inspection, we determined that about half of the reads carried a 3bp in-frame deletion, potentially explaining the green phenotype of plant #16. Yellow plants #1 (18% mutant reads for II) and #20 (53% mutant reads for II), transmitted respectively an inversion and a deletion allele of the gene region flanked by sites I and II. If these mutations were present in most cells, they may account for the unexpected observations since they would not amplify with either primer set used to prepare amplicons for Illumina sequencing. D. Correlation between mutation rate at the two sampled sites.

To assess the frequency and nature of the mutated sequences, genomic DNA was extracted from T1 leaves and the regions flanking the two guide RNA target sites were sequenced using Illumina technology. Sequence reads could be separated into two groups: reads containing the wild-type protospacer and reads containing a mutated protospacer. The leaf mutation efficiency was defined as the percentage of reads containing a mutated protospacer. The average leaf mutation efficiency across 31 T1 plants was 42% at the 5’ UTR target and 55% at the coding exon target. As expected, the plant phenotype had no obvious relationship with presence of a mutation at site *I*, but was well correlated to the presence of a mutation at site *II* (Fig. 2-C).

Next, we asked whether mutations at the two *CH1* sites were independent, i.e. whether high frequency of mutation at one site predicted the outcome at the other site. We found that mutation at site *I* accounted for ˜50% of the outcome at site *II* (Fig. 2-D) (Pearson r=0.48, p=0.0022), consistent with a specific cellular state, most likely expression of RC9, having a limiting effect on the probability of cutting. We concluded that both concurrent and single-cut instances were possible.

To further test the efficiency of the RC9 system with a different RNA expression strategy, three HOMEOBOX genes (*At5g66700* [*HB53*], *At4g36740* [*HB40*] and *At2g18550* [*HB21*]) were targeted by a single polycistronic guide RNA construct, consisting of three guides separated by tRNA motifs, driven by the U6-26 promoter ^13^. Similarly to *BRI1*, a plant transgenic for the sgRNAs was crossed to a plant expressing CAS9. Genomic DNA was extracted from leaves from 48 independent T1 individuals, the targeted regions were amplified and tested for the presence of mutated protospacers using restriction digests. The percentages of plants carrying at least one mutant allele were 33%, 67% and 60% for *HB21*, *HB40* and *HB53*, respectively. In other words, out of 48 plants, 9 carried at least one mutant allele for each of the three targeted genes.

Taken together, these three experiments using the RC9 system in various combinations all confirmed that the RPS5A promoter is an efficient promoter for producing high frequencies of somatic mutations.

### Mutations at non-genic sites

To test the efficiency of RC9 to target non-genic DNA, we selected three non-genic targets with multiplexed sgRNAs (Fig. 3-A, 3-B). For the 3 non-genic targets, the leaf mutation efficiency was 34.2, 96.8, and 99.1%, respectively (Table S2). In all three cases the most common mutation was a single base insertion between the third and fourth bases upstream of the PAM.

**Figure 3.**
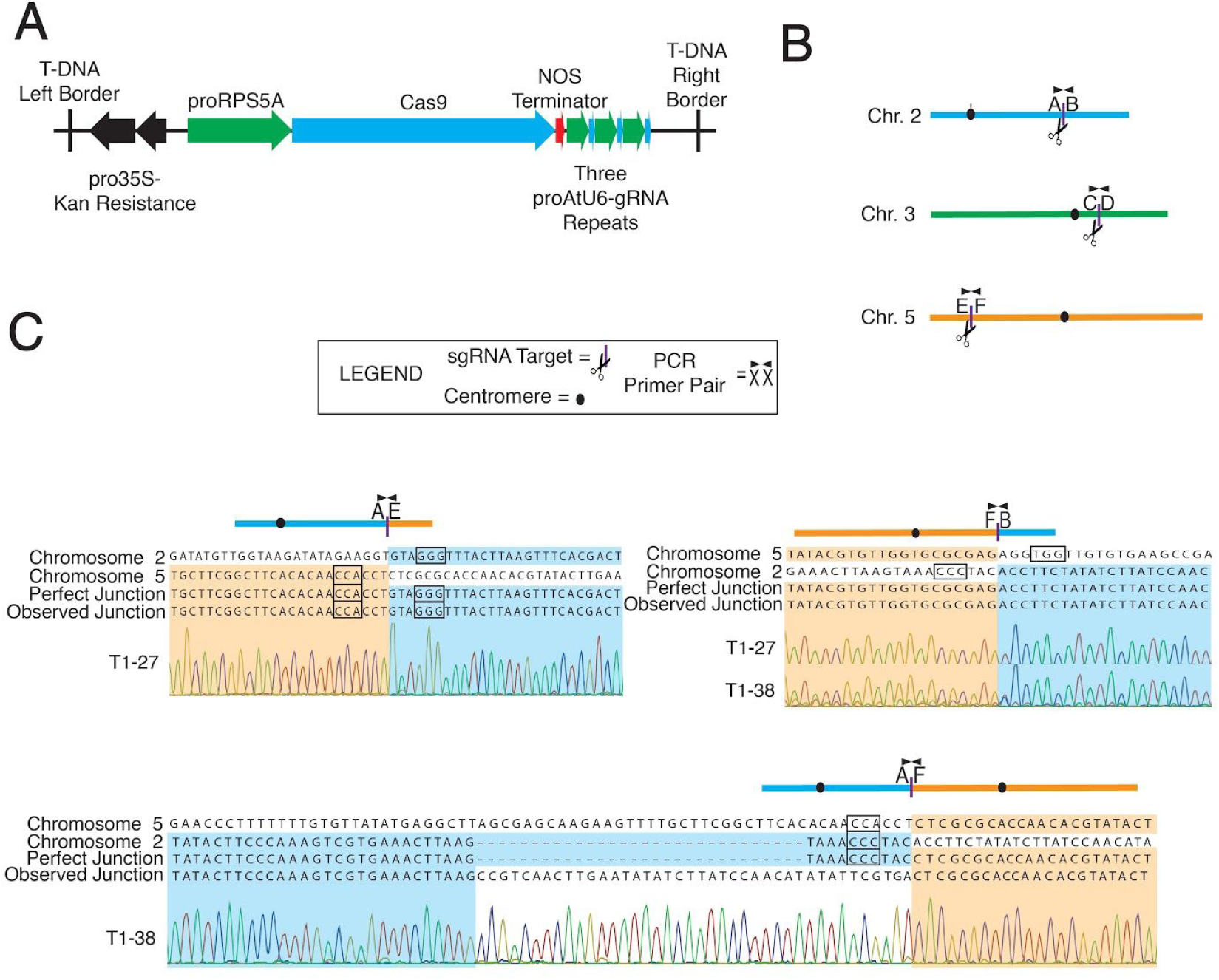
Cuts on different chromosomes can result in translocation. A. Structure of T-DNA expressing Cas9 and sgRNAs targeting intergenic sites NG1, NG2 and NG3 (Table S2). B. Location of intergenic targets on chromosomes 2, 3 and 5 with the corresponding PCR primers used to produce amplicons for Sanger sequencing. C. Alignment of T1 DNA Sanger sequences to in-silico translocation junctions show three perfect events and one containing both a deletion and insertion. PAM sequences are in black boxes.

To calculate the germline mutation efficiency, leaf DNA from T2 plants that are null segregants of the T-DNA were characterized. These sgRNA-RC9-negative T2 plants were Sanger sequenced at each target locus, and each individual was classified as containing 0%, 50% or 100% mutated copies at each non-genic target. This provides an estimate of the overall germline mutation efficiency, which was calculated by taking the germline mutation efficiency from each independent line and then averaging across independent lines (Table S3). The overall germline mutation efficiency ranged from 31.5-90.9% depending on the target (Table 1 and Table S2). While the sample size resulting from this method was small, the data show that germline mutations are common in randomly picked T1 plants, further demonstrating the effectiveness of RC9.

In summary, we observed good efficiency and early expression with RC9. Effectiveness of DNA editing, however, varied considerably, with two of the non-genic targets displaying the highest efficiency: nearly complete biallelic mutations in primary transformants in the germline and leaves.

### Multiplexed intragenic cuts can cause precise deletions and unexpected junctions

To investigate the frequency of *cis*-nonhomologous recombination following the targeting and cutting of different loci with CRISPR-Cas9, we first searched for deletions and inversions between the two intragenic *CH1* cut sites. The *CH1* cut sites are separated by 2,342 base pairs (Fig. 4-A). For practical reasons, we were unable to perform this test on T1 plants but were able to assess it in T2 plants. Leaf DNA of T2 plants were amplified with primers B and G that are upstream and downstream of the targets, respectively (Fig. 4-A). Appearance of a 519 bp band suggests presence of the deletion. The percentage of all analyzed T2 plants containing at least one copy of the deletion ranged from 0-95% depending on the T1 parent (Table S4). Twenty-three of the 164 T2 plants that contain deletions were Cas9-negative, indicating that the deletion had been transmitted meiotically from a parent. The twenty-three Cas9-negative alleles were found amongst 7/32 T2 families, suggesting frequent formation of deletion alleles in the germline.

**Figure 4.**
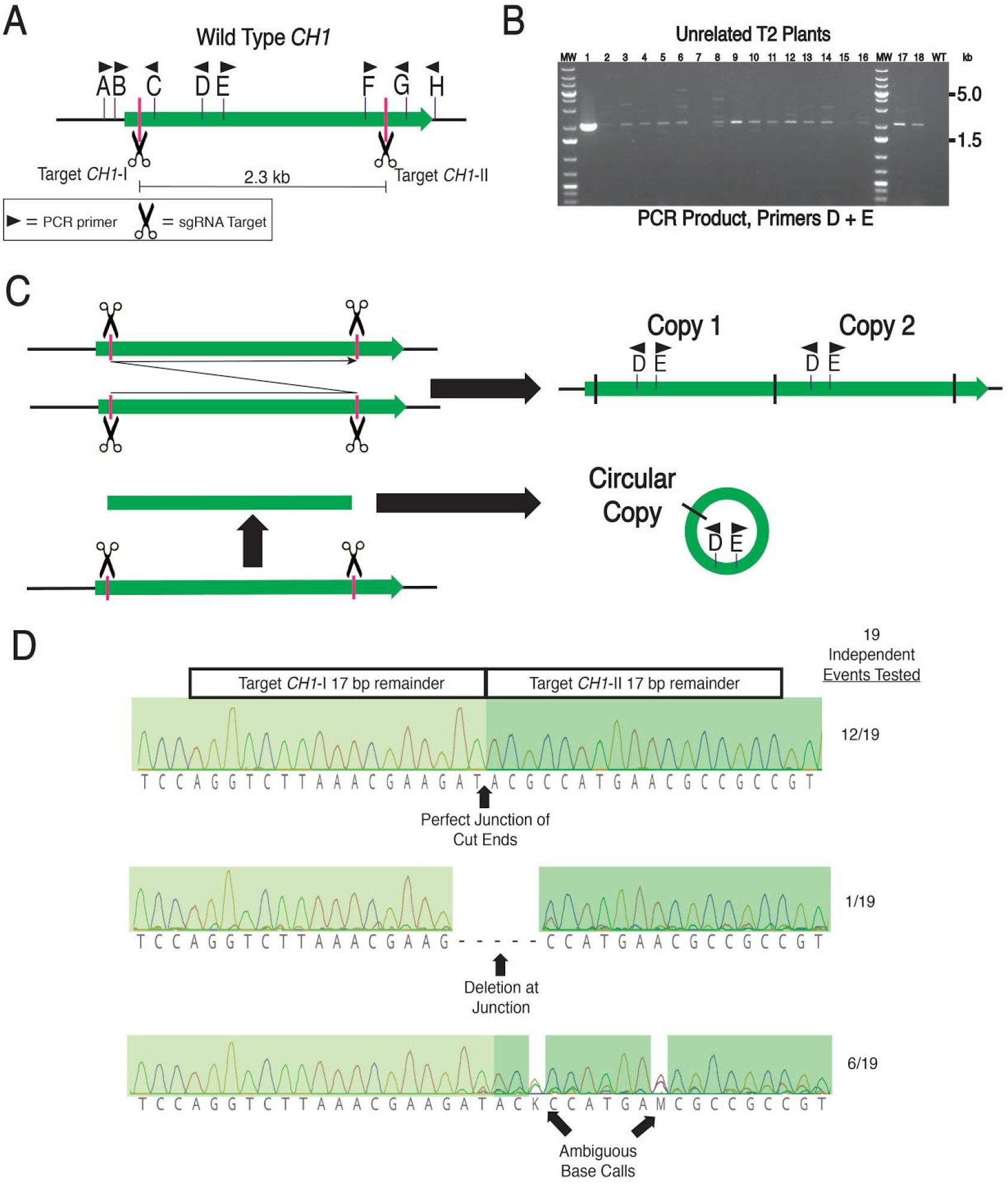
Dual cuts in *CH1* result in perfect recombination junctions suggestive of duplication or circularization. A. Map of the *CH1* gene (*At1G44446*) showing the position of the primers used in this study (arrows). B. When PCR primers D and E are used together, bands are regularly found. Each column used genomic DNA from a single leaf of a single T2 plant from eighteen different primary transformants containing proRPS5a-Cas9 with sgRNAs for the two CH1 targets. Column #1 is from an individual with a confirmed duplication that is present throughout the germline and plant, which is why it is the brightest band (See Fig. 5). The far right lane corresponds to the wild-type control and yields no PCR product, as expected. MW = Molecular Weight ladder. C. These recombinant bands are consistent with either a duplication or circularization of the segment between the two CH1 cuts. D. When using PCR primers C and F (see Fig. 3-A), recombinant duplication or circularization junctions were detected in the progeny of eighteen different primary transformants out of thirty two primary transformants. PCR bands from one plant from each of the eighteen families were Sanger sequenced. Twelve of the junctions are perfect, meaning that no bases were added or subtracted from the exact CRISPR cut sites. One junction exhibited a 5 base deletion. Six of the junctions have ambiguous base calls.

There is the possibility that an excised fragment will be reintegrated in the opposite orientation, resulting in an inversion of the 2.3 kb segment. To search for these events, we used a primer pair that should only amplify an inverted junction (Fig. 4-A). Inversion bands were found in 24/82 T2 plants from 5/13 T1 parents that were screened for inversion junctions (Table S4). Eight of those 24 T2 plants were checked for the inversion junction at the other side of the *CH1* segment using primer pair C + G (Fig. 4-A). Results from 7/8 plants were consistent with a full inversion, with expected inversion junction sequences on both sides of the segment (Table S4).

Theoretically, another type of recombination junction could result from either segment duplication or circularization (Fig. 4-C). We searched for these events next, using primers that would only produce a band if the corresponding “inner” junction is present (Fig. 4-B, 4-C). We tested 291 T2 plants derived from 32 primary transformants finding a total of 81 positive T2 individuals in 18 families. More than half of the T2s in 7 families contained this kind of recombinant junction, suggesting that it forms at high frequency. Amplicons from 19 different events were digested with XbaI, and the results matched the expected recombinant junction. One band from each of the 19 junction-positive primary transformants was Sanger sequenced. Twelve of the sequences matched the predicted, perfect junction (Fig. 4-D). The remaining seven sequences contained at least one mutation in the residual protospacers at the junction (Fig. 4-D).

### Duplication of a 2.3kb segment

The presence of the inner junction (above) suggested either duplication or circularization of the DNA segment flanked by the CRISPR cuts. Testing for circles failed to provide reliable evidence for their existence (see Methods). For this reason, we pursued the hypothesis of duplication. To probe for potential duplications, we used primers B and G (Fig. 5). From wild-type control templates, these primers amplified a 2.8kb DNA fragment containing the 2.3 kb segment of interest (n=35). Using these primers on a template consisting of the direct duplication of the 2.3 kb fragment is expected to result in a 5.2 kb PCR product. We screened an average of 9 T2s from 32 *CH1*-CRISPR families. In 88 T2s from 18 families, we amplified concurrently a predominant wild-type product (2.3 kb segment in a 2.8 kb PCR band) and a weaker duplication product (4.6 kb segment in a 5.2 kb PCR band). We reasoned that the two-product pattern resulted from presence in the template of both allele types, wild-type and duplicated, perhaps resulting from heterozygosity or from chimeric individuals. Spurred by this finding, we searched for individuals where the duplication allele was fixed.

**Figure 5.**
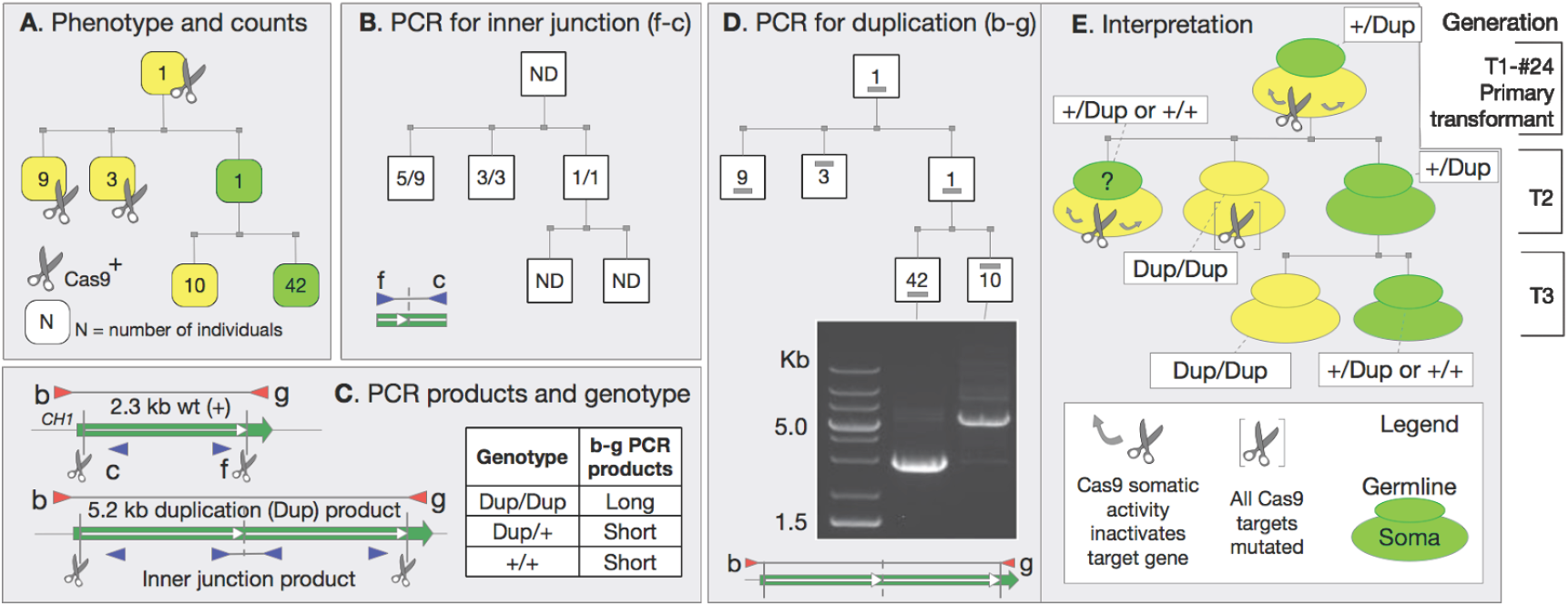
Induced duplication in *CH1.* The same pedigree starting with yellow T1#24 is repeated in A,B,D, and E. Each box in T2 and T3 generation represents a class of individuals whose numbers are noted inside the box. A. A single green, Cas9-free T2 carries a knockout allele. B. Analysis of “inner” junction PCR product (Fig. 4-C). C. *CH1* gene map with cut sites and primers used for PCR. The table illustrates how in the genotyping assay the long duplication PCR product is not detected,probably because of differential amplification efficiency. D. Genotyping assay for the long PCR product found in duplication homozygotes. For a precise tandem duplication of the 2.3 kb segment, the expected PCR band sizes are 2.8 kb for single copy and 5.2 kb for duplication. (C). E. Soma and germline genotypes consistent with the observations. The yellow progenitor T1-#24 was evidently chimeric suggesting multiple mutagenesis in soma leading to virtually complete inactivation of *CH1* and yellow phenotype. Its germinal cells were heterozygous, carrying a wild-type *CH1* allele and an inactive *ch1* duplication allele. The T2 generation displayed a 3:1 ratio of large to small PCR products (Chi-squared p-value = 0.337). Yellow T2s displaying the large PCR product are likely homozygotes that inherited the inactive *ch1*-duplication allele. The T2s that displayed the small PCR product inherited at least one wild-type *CH1* allele. Cas9-positive T2s were all yellow and fall in two categories: 9 of them display the small PCR product. The 4 T2s in this group of 9 that did not display the junction fragment inherited two wild-type alleles. Their yellow phenotype is consistent with virtually complete mutagenesis of their soma. The 3 T9 that display the large duplication product were most likely homozygous for the knockout duplication allele and not chimeric. The single Cas9-negative T2 was green and must have been heterozygous because it formed T3 progeny of two phenotypic classes. Ratios of large:small PCR products (B), and yellow:green pigment individuals (C) coincided (42:10), fitting a mendelian 3:1 F2 ratio (Chi-squared p-value = 0.873). The Cas9-free branch provides good evidence for the model in E.

The analysis of plant family #24 provided both evidence for duplication and an interesting example of inheritance after CRISPR-Cas9 transformation (Fig. 5). The family was founded by T1 (primary transformant) individual #24, which was yellow and positive for the duplication junction (Fig. 5-A). Twelve of 13 T2s were similarly yellow, but also Cas9 positive (Fig. 5-A). A single Cas9-negative T2 was green, consistent with chimerism in T1 #24. Upon selfing, this single green T2 plant produced 42 green and 10 yellow T3 plants, a good fit for the 3:1 ratio expected from a heterozygote (P of chi square = 0.337). PCR amplification of the *CH1* segment with flanking primers using DNA template from yellow T3 plants displayed the duplication product (Fig 5-D). Green T3 plants yielded either a pure wild-type product or the two-product pattern. We interpret these results as follows: the primary T1 contained at least two alleles: the wild-type and a duplication allele and transmitted both to its progeny (Fig. 5-C). The T2s appear to result from segregation of a heterozygous T1 germline, but their phenotype is confounded by independent assortment of the Cas9 gene. In the presence of Cas9, heterozygous T2 that would normally be green are mutated in the soma and appear yellow. The single Cas9-negative T2 plant, was green and heterozygous, having a wild-type allele and a duplicated mutant allele. T3s produced by this plant had phenotype consistent with their genotype (Fig. 5 and Table S4). Heterozygous T3 progeny were green and preferentially yielded the smaller, wild-type PCR allele. Plants homozygous for the duplicated allele were always yellow and yielded the large, duplication PCR product only (Fig. 5A, Fig. 5D). In conclusion: activity of RC9 in the soma can produce an apparently fully mutant phenotype; reliable analysis is only possible in Cas9-free segregants; PCR products from heterozygotes are skewed toward the shorter product; and duplication of the *CH1* segment results in a defective allele.

The recombinant junction product found in progeny of 18 primary transformants could have multiple causes. It could result from the presence of chimeric and sub-stoichiometric (on a whole genome scale) template corresponding to duplication or circle products.

### Duplication of a 8.5kb segment

Next, we tested the effects of cleaving a larger segment. We targeted two sites 8.5 kb apart that delimit a segment containing three genes, including LEAFY COTYLEDON 1 (*LEC1;* At1G21970). Forty-three primary transformants were tested for the duplication junction using LEC1 primers A and B (Fig. 6-A). Eighteen of the 43 PCR reactions produced a band of the expected size. Sixteen of the bands were confirmed by Sanger sequencing to be the duplication (or, potentially, circularization) junction. Seven of the 16 chromatograms suggested perfect junctions, and 9 of the chromatograms indicated indels in the junction (Fig. 6-B). Together with the CH1 analysis above, this indicates that Cas9-driven cutting of targets 2 to 8 kb apart can result in tandem duplications. The potential for circle formation remains to be investigated.

**Figure 6.**
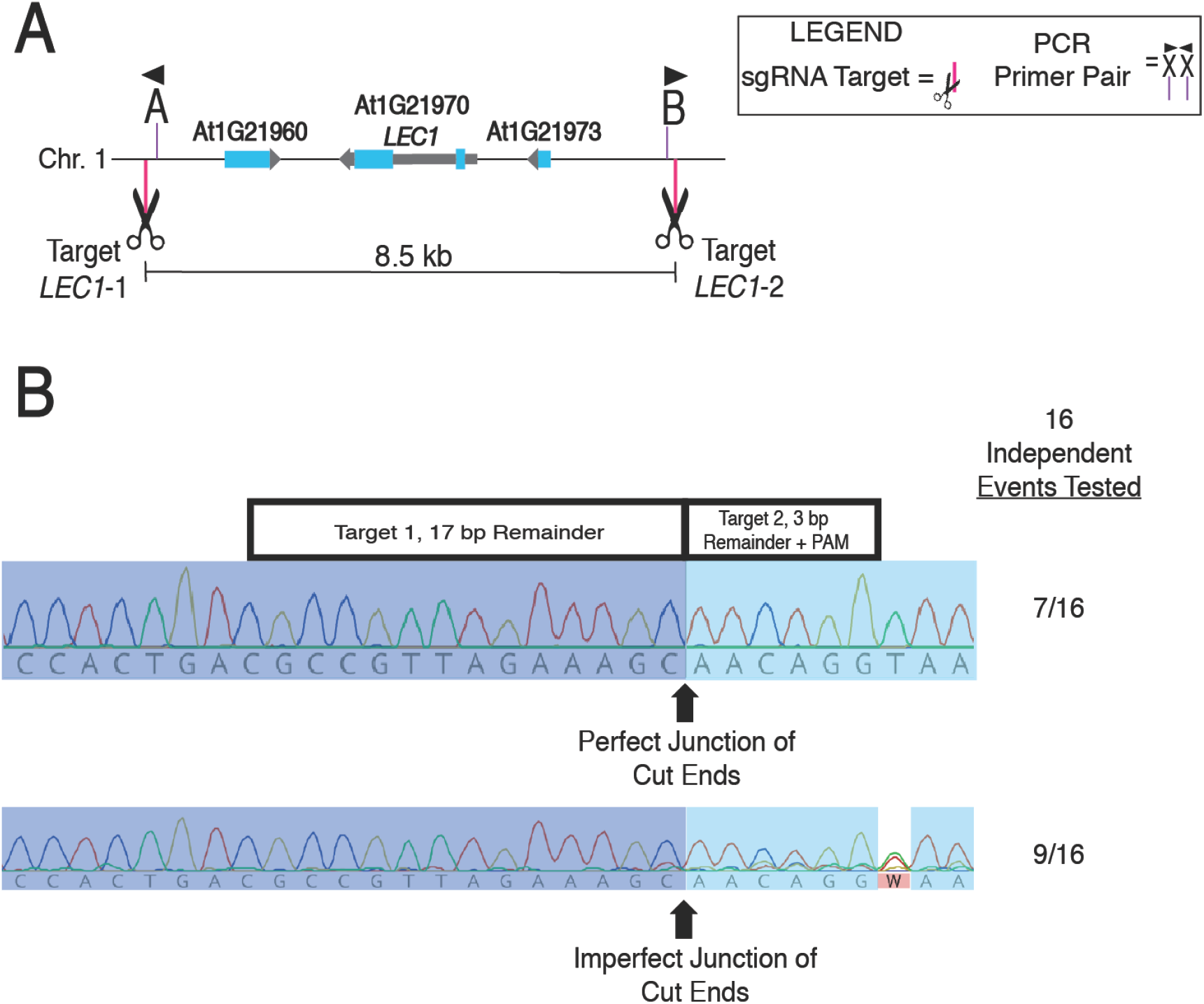
Dual cuts flanking the entire *LEC1* gene frequently results in perfect recombination junctions consistent with circularization or duplication. A. Map of the locations of the Cas9 target sequences flanking an 8.5 kb segment containing two genes and one pseudogene. B. Eighteen out of 43 primary transformants yielded PCR bands using primers A and B that were of the expected size for segment duplication or circularization. Sequencing with Sanger technology revealed that sixteen bands were the expected recombinant junctions; the other two bands were not sequenced. Each of the sixteen chromatograms is independent from the others because each band came from an independent primary transformant. Seven of the 16 junctions were perfect and 9 contained indels or chimerism near the junction.

### Induced dsDNA breaks at nonhomologous sites on different chromosomes can cause translocations with precise junctions

When targeting more than one genomic locus, it is possible that simultaneous cleavage can result in interchromosomal translocations between nonhomologous loci. To detect these events, we designed PCR primers assaying interchromosomal exchange between the three unrelated non-genic targets: NG1, NG2 and NG3 (Table S2, Fig. 2). In this assay, amplification using primers from different target chromosomes is only expected if a translocation occurred. Up to 34 T1 individuals were tested with each possible primer pair (Table S5). PCR amplification using flanking primers were successful from two T1 plants. PCR products in one of these two plants suggested a Chr.2/Chr.5 translocation as well as the reciprocal translocation resulting in two monocentric recombinant chromosomes (Fig. 2-C). The other translocation contained a Chr.2/Chr.5 band that suggests the presence of a dicentric chromosome (A/F junction in Fig. 2-C). Nineteen progeny of the plant containing reciprocal F/B and A/E translocations were tested for presence of the F/B junction. Ten progeny of the plant containing a single F/B junction were tested. None of the tested progeny contained a transmitted F/B junction.

In order to further characterize these events, the junctions of the four non-genic translocations were sequenced using Sanger technology. All three monocentric translocations exhibited perfect junctions between the two non-homologous sequences, with no insertions or deletions (Fig. 2-C). The junction of the presumed dicentric recombinant chromosome contained a 10 base deletion and an unknown (no BLAST hits) 40 base insertion (Fig. 2C).

We also used 9 primer pair combinations to screen for translocations in 39 F1 plants in which three HOMEOBOX genes were targeted. No translocations were found between the 3 HOMEOBOX gene target sites (Table S6).

### Targeting two loci in one gene and one locus on a different chromosome frequently causes translocations with precise junctions

To further test if cutting targets on different chromosomes can result in translocation, we targeted loci on different chromosomes, with two cuts separated by 1.7-2.0 kb and one cut on a different chromosome (Fig. 7A, 7B). The two targets separated by 1.7-2.0 kb were in the promoter region of the genes *PIGMENT DEFECTIVE 334* (*PDE334*; At4G32260) or *RAB GTPASE HOMOLOG E1B* (*RABE1B*; At4G20360) (Fig. 7C). The target on another chromosome was near the transcription start site (TSS) either *LEC1* or *WUSCHEL 1* (*WUS1*; *AT2G17950*). Induced nonhomologous recombination between the targets was detected by PCR of primary transformant (T1 generation) DNA using primers that would allow amplification of the predicted junctions (Table 1).

**Figure 7.**
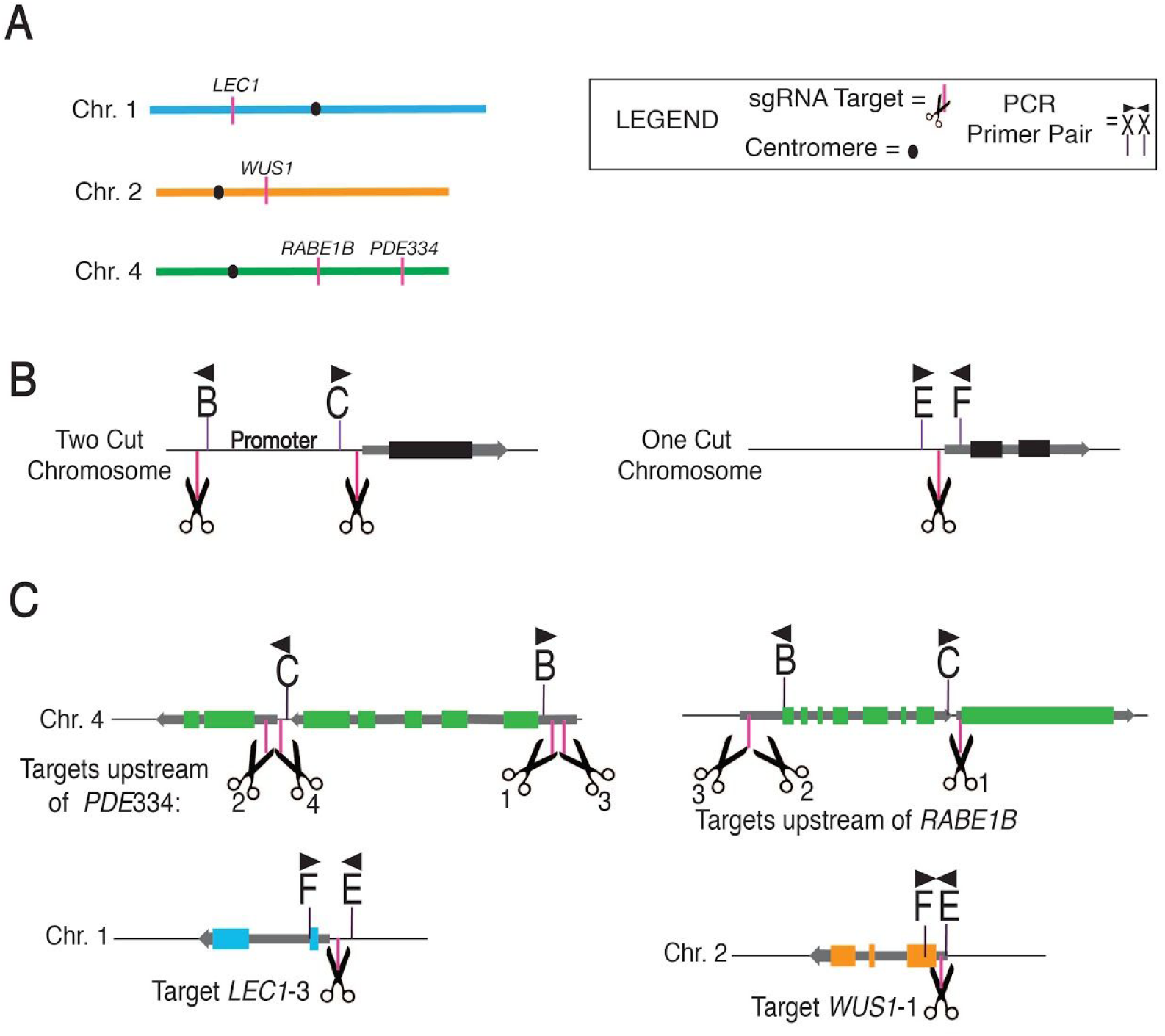
A. Physical location of CRISPR-Cas9 targets on *A. thaliana* chromosomes. B. Generic model for positions of cuts and PCR primers when two cis cuts and one trans cut are made. The primers correspond with those of Table 1. C. Location of targets in the 5’UTR and promoter of *PDE334*, *RABE1B*, *LEC1* and *WUS1*. In this experiment each construct contains 3 sgRNAs: one targeting the 5’ and one targeting the 3’ end of either the *PDE334* or *RABE1B* promoter and one targeting near the TSS of *LEC1* or *WUS1*.

One combination of targets that frequently induced translocations was when two cuts were made 1.8 kb apart on Chromosome 4 upstream of *PDE334*, and the third cut was made 8 bp upstream of the *LEC1* TSS (Fig. 8-A, 8-B) ^*41*^. Using a primer near the 3’ end of the *PDE334* promoter and a primer in the *LEC1* gene, faint but clear PCR bands of the expected size were found in 36/121 primary transformants (Table 1., Fig. 8-C). Sanger sequencing 20 of the 37 PCR products confirmed that they represented the expected junction and revealed 10 translocation alleles, i.e. having different sequences. The perfect junction allele was predominant in 4 of the 20 chromatograms (Fig. 8E, 8-F, Fig. S2). A very similar result was obtained in a different set of primary transformants, in which a smaller fragment containing the *proPDE334* was targeted by moving the transcription start proximal guide 95 base pairs upstream of the guide from Fig. 8. A PCR screen of genomic DNA from a single leaf of each of 163 primary transformants revealed that 25/163 primary transformants contained the translocation (Table 1, Fig. 7-C, Fig. S3). Moving the *proPDE334* 5’ guide upstream by 40 bp let to a frequency of 37/109 primary transformants containing the 3’ *proPDE334*-*LEC1* junction (Table 1, Fig. 7-C, Fig. S4). When the 5’ end of *WUS1* was targeted instead of *LEC1*, 22/74 primary transformants were found to contain the pro*PDE334*-pro*WUS1* translocation junction in which the *PDE334* promoter and *WUS1* promoter should produce convergent transcription (Table 1, Fig. 7-C, Fig. S5).

**Figure 8.**
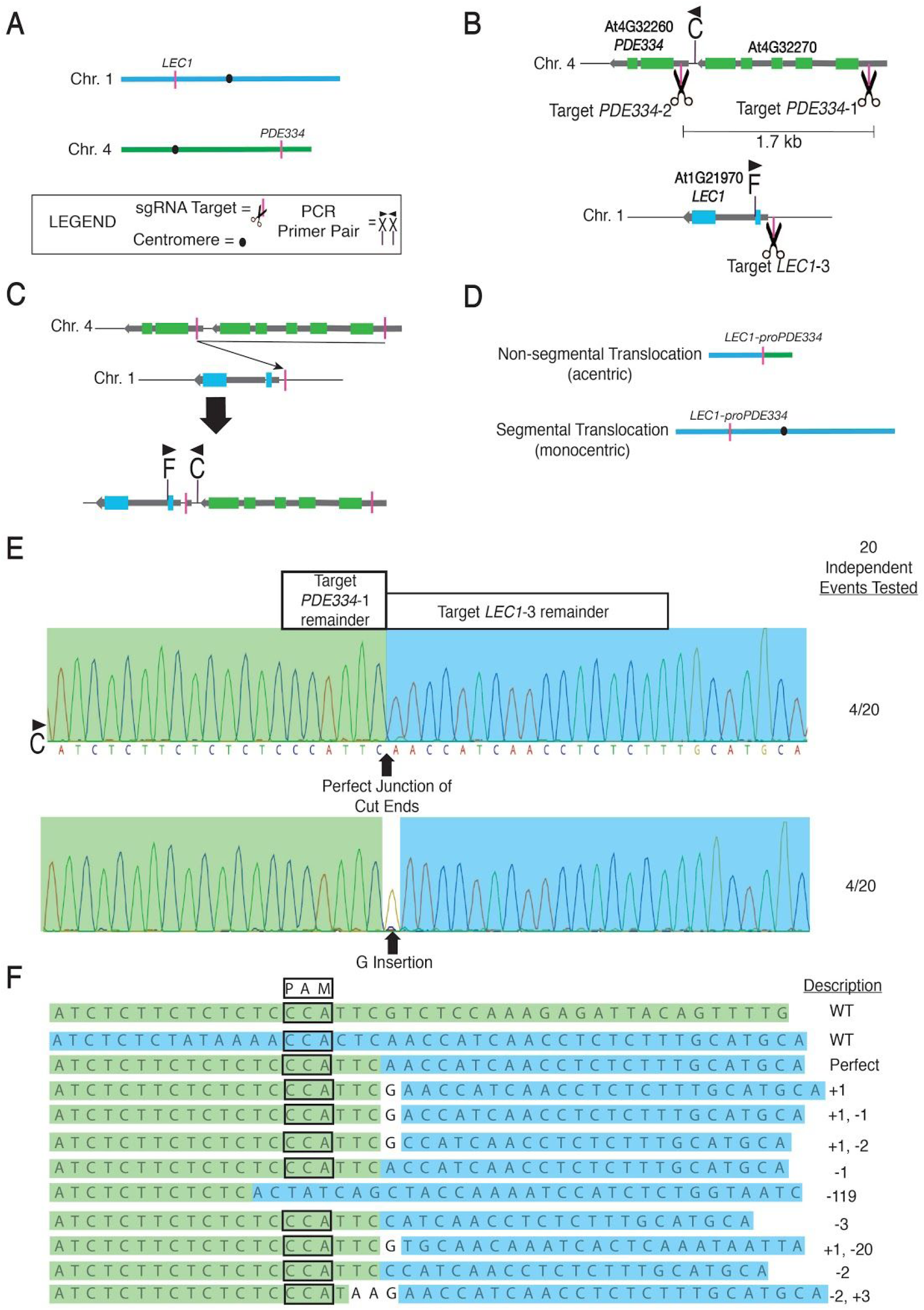
Frequent formation of a precise nonhomologous translocation. A. Map of the locations of At1G21970 (*LEC1*) Chromosome 1 and At4G32260 (*PDE334*) on Chromosome 4. B. Map of the locations of the Cas9 target sequences estimated to flank the *PDE334* promoter and the end of the *LEC1* promoter, for one construct (See Table 1). C. Using PCR primers PDE334 C and LEC1 F, translocation junctions were found in 36/121 T1 plants. D. The C/F junction may be due to a segmental or non-segmental translocation. In this case, a non-segmental translocation would result in formation of an acentric chromosome. Segmental translocations of this size are expected to always result in a monocentric chromosome. E. Twenty of the 36 translocation bands were sequenced with Sanger technology using the C primer. Each of the 20 chromatograms is independent from the others because each band came from an independent primary transformant. Four of the 20 chromatograms exhibit perfect junctions and no mixed peaks. F. In addition to the perfect junction, 9 additional alleles were found in the 20 chromatograms. Indel sizes are described as compared to the perfect translocation junction.

As an attempt to control translocation orientation, some perfect translocation junctions were designed to reform a target sequence that would be re-cut, thus hypothetically disfavoring fixation of that junction (Table 1, Fig. S6). Similarly, some predicted perfect circles were designed so that they would be cut open at the circle junction, thus possibly increasing the time spent as a linear segment (Table 1, Fig. S6). However, neither of these approaches made a significant difference in the orientation or frequencies of translocations (Chi Square p = 0.791 for re-cut translocations and p = 0.1065 for re-cut circles affecting the frequency of translocations). A limitation is that the experiment depends on formation of perfect junctions.

Some fusions might overexpress *LEC1* and *WUS1*, genes that induce ectopic embryos ^42,43^ (Table 1). The resulting transgenics, however, appeared normal. As shown in Fig. 9, mitotic stability of these promoter-gene fusions would require segmental translocation of the promoter fragment and repair of all cut sites. A chromosomal translocation involving two chromosomal ends, on the other hand, should form acentric fragments (Fig. 9). To investigate the possibility of segmental transfer of the PDE334 promoter, we tested PCR amplification of the upstream PDE334 junction to the upstream LEC1 DNA (Fig. 8), but this failed in all but 2 of the 121 tested plant from which progeny was not retrieved (Table 1).

**Figure 9.**
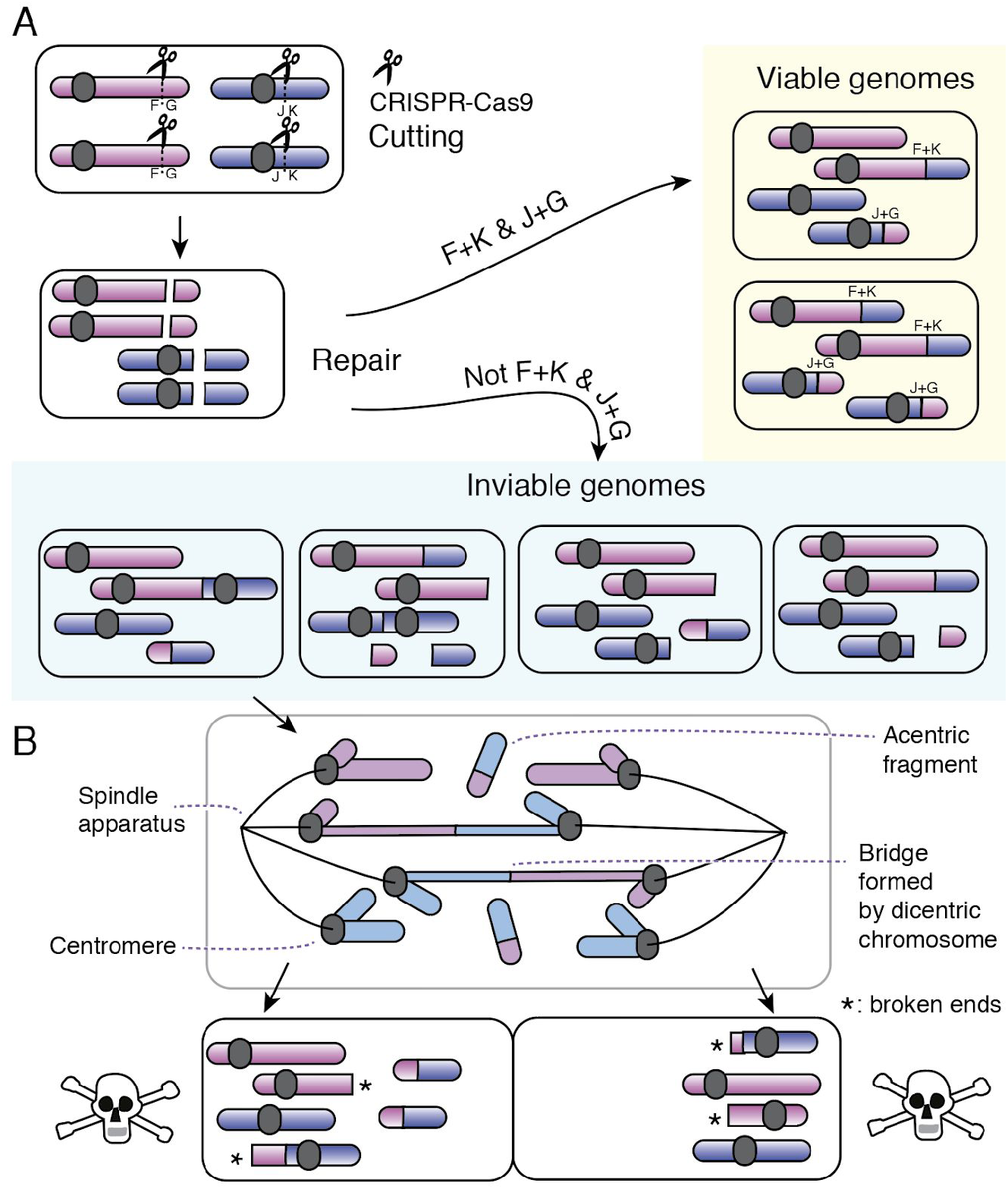
Viable and inviable repair events after occurrence of two dsDNA breaks. A. Two targeted dsDNA breaks are introduced in two pairs of chromosomes, red and blue. When these cuts are repaired by events other than reversion, only the reciprocal translocation event illustrated as “F+K & J+G” will result in viable cells. Meiosis is predicted to yield 50% and 100% viable gametes from, respectively, top and bottom cell. B. Other translocation junctions (“not F+K & J+G” path), or end healing, as exemplified for the leftmost inviable cell, are predicted to be deleterious or lethal because acentric or dicentric fragments cannot be inherited regularly by daughter cells. Anaphase and karyotypes of the daughter cells are displayed. The case illustrated assumes that target chromosomes are cut at least once. Partial cutting and fusions of the type illustrated would result in segmental aneuploidy with the connected deleterious effects.

Altogether, in 881 tested T1 plants, we found 195 instances of positive signals. We sequenced 91 of the corresponding PCR products and identified the diagnostic sequence in 84 of these (Table 1, Fig. S2, Fig. S3, Fig. S4, Fig. S5). We concluded that interchromosomal fusion is possible for a majority of the tested cut ends.

## Discussion

The *RPS5A* and *U6-26* promoters were used to express Cas9 and multiplexed sgRNAs, respectively, early in development to achieve efficient mutagenesis in somatic and germline tissues. We confirmed efficiencies similar to those previously reported ^10^. In addition to the formation of deletions between close CRISPR sites, we demonstrated the formation of chromosomal rearrangements of considerable interest: translocations and duplications that are often religated perfectly.

### Somatic vs germline mutation

The recurrence of a whole plant phenotype is insufficient proof that a mutation occurred in the germline ^17^, and Cas9-free segregants are required to determine germline inheritance. This is exemplified by family *ch1-24* (Fig. 5). While uniformly yellow, the founder T1 must have been chimeric because it transmitted both a functional allele and a knockout allele with a tandem duplication event. The mutant phenotype displayed by its body was most likely produced from developmentally late mutations. Consistent with the high frequency of these hypothetical somatic events, all Cas9-positive T2 plants were yellow. This despite genotyping evidence (short PCR allele, Fig. 5) that this class, which represented 75% of the progeny, inherited the wild-type allele and should be green. We infer that Cas9-induced knockout mutations of the wild-type allele were preponderant in the soma. The dominant action of the wild-type allele was only manifest in the single Cas9-negative T2, which was green. The progeny of this T2, in turn, followed the Mendelian pattern expected from a heterozygous parent. Around 1/5 of the T3 were yellow and positive for the inactivated allele. The rest of the T3 were green. In conclusion, in the presence of active Cas9 and sgRNA, somatic mutations are common and often confer the recessive phenotype. Establishment and germline transmission of mutant alleles likely depends on CRISPR-Cas9 activity in meristematic cells, such as is the case for RPS5A.

### Duplication

Our data indicate that duplication of a segment flanked by two target sites is possible. Duplication of a 2.3kb region was transmitted meiotically in one line (Fig. 5). Duplication could also be found at a second independent 8.5 kb segment containing three genes. In a limited set of 8 tested progeny we could not document transmission. The mechanism of the segmental duplication may be explained by a simple nonhomologous translocation, similar to that represented in Figure 8 and Table 1 except occurring between sister chromatids or homologs. A translocation between sister chromatids could explain the duplication in individual #24 (Fig. 5) because we found no deletion allele in any of the 52 tested progeny. A deletion allele could persist in the same nucleus if recombination was between homologs (depending on cell cycle phase), while a deletion allele formed in a sister chromatid will be always be transmitted to a different daughter cell (Fig. S7). Duplications were detected by amplifying a novel junction. This assay, however, will also detect a circle (Fig. 4-C), and amplification of the large diagnostic fragment that spans the duplication is problematic in a heterozygote (Fig. 5-D). Therefore, we cannot rule out that formation of a circle may contribute to the junction signal. In any case, a tandem duplication itself can engender circles ^44,45^ further complicating the analysis. The targeted regions lacked known replication signals and these circles would be transient. Our efforts to detect genomic circles by DNA exonuclease or phi29 polymerase enrichment yielded erratic results (see Methods). In conclusion, we demonstrate that duplications are formed.

### Translocation

Translocation requires that cut chromosomal ends meet in the nucleus before a different repair reaction occurs ^46^. Plant nuclei are large and it is not clear that cut chromosomal ends have sufficient mobility to meet efficiently. Surprisingly, we detected 195 interchromosomal targeted translocations in one or two leaf samples of 881 tested primary transformants. Some of these could have resulted in promoter fusion and overexpression of *LEC1* and *WUS* coding regions leading to developmental abnormalities ^42^. The absence of such symptoms in our experiment could have at least two explanations. First, fusions that do not involve segmental transfer generate chromosomal translocations that produce unstable acentric and dicentric chromosomes (Fig. 9). Cells carrying acentric or dicentric events are unlikely to proliferate. Second, it is possible that these fusions are not sufficient to trigger overexpression or if they do, they have deleterious consequences. Exploitation of interchromosomal fusions to engineer chromosomes or alter gene expression will benefit from careful planning to ensure neutrality for efficient transmission. While these recombination products are likely to be useful, they could occur as undesirable byproducts of the experimental aims, potentially triggering genome instability.

### Conclusions

In this study we examined the effect of expressing Cas9 with a promoter active in early embryo and in meristematic tissues. By cutting at different chromosomal sites, we document the formation of precise junctions forming duplications and translocations. Mechanisms resulting in these products, the precise conditions favoring their formation, and the generality of these responses remain to be determined. Harnessing the efficient formation of precise junctions between non-homologous DNA ends could lead to multiple useful applications.

## Materials and Methods

### Fluorescent Microscopy

To visualize centromeric GFP and whole nuclear tdTomato fluorescence signals, young embryos were dissected under a stereomicroscope from unfixed *Arabidopsis* ovules (42-45 hrs after pollination) immersed in 0.1X PBS using a pair of fine tungsten needles (Cat# 10130-05 from Fine Science Tools-USA). Dissected embryos were collected with a small volume of 0.1X PBS and transferred into 5ul of mounting medium (1ug/ml DAPI, 50% glycerol and 0.1X PBS) on glass slides using a fine glass pipette and gently covered with a thin coverslip (No.1.5, 22×22mm). Samples were imaged within a few hours after dissection using a Deltavision deconvolution microscope with Z-stack capability. Images were captured using 60X lens with an exposure time of 0.2-0.3 seconds and 0.2 to 0.5 microns optical sections. Raw data (Z-stacks from each embryo) were deconvoluted using Softworx software (Applied Precision/GE Healthcare). Deconvoluted Z-stacks (3D-image) were analyzed and converted into 2D projections using the Imaris software and exported as a TIF files for further editing using Adobe Photoshop and Illustrator software.

### Creation of proRPS5A::Cas9

The ORF of Cas9 gene from plasmid pDE-Cas9 ^6^ was amplified with primers Cas9HP_fwd and Cas9HP_rev (Table S7). The RPS5a promoter was amplified from *Arabidopsis thaliana* Col-0 genomic DNA with primers RPS5aPro_fwd and RPS5aPro_rev (Table S7). These two fragments and Hpa I-digested pPLV02 (De Rybel et al 2011) were assembled using Gibson Assembly Master Mix (NEB) to make plasmid pPLV02_proRPS5a::Cas9.

### Creation of non-genic triple sgRNAs

Protospacers for three non-genic targets (Table S1, S8) were each inserted into pEn-Chimera vectors at the sgRNA site (Fauser et al., 2014). The proU6-26-sgRNA DNA was amplified with unique ends added for Gibson Assembly. Plasmid pPLV02_proRPS5a::Cas9 was cut with Xbai and then was ligated to three proU6-26-sgRNA fragments using Gibson Assembly Master Mix (NEB).

### Creation of 3-Fragment Multisite Pro system

The palindromic BbsI sites for sgRNA cloning into pEn-Chimera were replaced with BsaI sites for pDONR compatibility. The U6 promoter, sgRNA scaffold, and S. pyogenes terminator were amplified from pEn-Chimera (Fauser 2014) with the addition of attB1/4, attB4r/5r, and attB5/2 tags and respectively recombined with pDONR221 #1, pDONR221 #2, and pDONR221 #3. This produced the entry clones pEn_Comaira.1, pEn_Comaira.2, and pEn_Comaira.3. The NOS promoter, KanR CDS, and NOS terminator were amplified from pBI 121 (Invitrogen) with attB1/4 tags and recombined with pDONR221 #1 to make pEn_nos:kan.1.

The RPS5a:Cas9 cassette was amplified with attB5/2 tags and recombined into pDONR221 #3 to produce pEn_RC9.3. All PCR was performed with Phusion HF master mix. All BP reactions were performed with BP Clonase II (ThermoFisher).

### Synthetic CRISPR-Cas9 activation

For targeting *BRI1*, three independent lines transformed with *proRPS5a::Cas9* were pollinated with seven independent lines transformed with the *BRI1*-targeting sgRNA (Table S8). Seedlings were obtained from 16 of the 21 possible combinations and these families were scored for the *bri1* dwarf phenotype. Line CAS9-4 (Table S8), which was the best of the three CAS9 transformants tested in *BRI1* inactivation, was used for *VRS* gene inactivation by crossing to the sgRNA line.

### Targeting of CH1 and the 8.5 kb segment containing LEC1

Design for the two sgRNA sites on *At1g44446* (*CH1*) was performed by analyzing the protein coding sequences on exons 1 and 9 of the *CH1* gene that contain PAM sequences. These sequences were verified using BLASTn to ensure they are unique targets and are provided in Table S1, along with the oligos used cloning in Table S7. In order to utilize the the 3-Fragment Multisite Pro system, we generated pEN-Comaira.ch1.1 and pEN-Comaira.ch1.2 by integrating double-stranded sgRNA oligo sequences with compatible overhangs (Table S7) into purified, BsaI digested, pEN-Comaira.1 and pEN-Comaira.2 respectively. All entry clones were verified by sequencing. The final reaction was performed by combining 50 ng of pEN-Comaira.ch1.1, pEN-Comaira.ch1.2 and pEN-RC9.3 and pEARLEYGATE202v2 with the recommended reaction volume using LR Clonase II plus (Invitrogen Catalog No. 12538120). The final construct containing all three inserted fragments was verified by Sanger sequencing and electroporated into Agrobacterium tumefaciens GV3101 for transformation of wild-type Col-0 by floral dip. The same process was done for the construct containing two sgRNA sequences targeting an 8.5 kb segment around At1g21970 (*LEC1*). The primary transformants were selected with glufosinate.

### Targeting of the PDE334 and RABE1B promoters and near the TSS of LEC1 and WUS1

Double-stranded oligo sequences of the sgRNA guide sequences for *PDE334* Target 1, *PDE334* Target 2 and *LEC1* Target 3 were ligated into pEN-Comaira.1, pEN-Comaira.2 and pEN-Comaira.3 respectively. The Gateway destination vector pEARLEYGATE202v2 was modified by inserting the same proRPS5A-Cas9 sequence used throughout this manuscript to make destination vector pDE-RC9. The multisite Gateway reaction, transformation methods and selection was performed as in the *CH1* experiment. The other combinations of *PDE334* or *RABE1B* with *LEC1* or *WUS1* guides were also constructed this way.

### Floral Dip

*Agrobacterium tumefaciens* GV3101 was transformed by electroporation for each complete construct. Wild Type *Arabidopsis thaliana* inflorescences were dipped into a solution containing *Agrobacterium tumefaciens* GV3101 that inserts the T-DNA into the egg cell genome. All plants were transformed using the floral dip method ^47^.

### Ampliseq design and analysis

Illumina sequencing of amplicons spanning the CRISPR cuts sites was performed as previously described ^17^. A Python script was used for quality control of the reads[IH1]. The number of reads for each unique sequence were counted. Only reads representing the top 100 most common sequences were used in the following analysis. All the top 100 sequences were visually compared in Geneious as a further check that they are of the target loci. Reads with an intact protospacer are grouped together as “Non-Mutated”. Reads with a mutated or deleted protospacer were grouped together as “Mutated”. The somatic (non-germline) mutation efficiency for each target was defined as (# Mutated Reads)/(# Mutated Reads + Number Non-Mutated Reads)X100%.

### Breakpoint PCR for Nonhomologous Recombination Junction Detection

Rosette leaves were ground in a CTAB solution to extract genomic DNA. PCR primers were designed that should only yield PCR products of the expected size if a specific recombination event had occurred. PCR was used to detect nonhomologous recombination events including translocations. Junctions were confirmed with DNA sequencing by Sanger technology.

### Circle enrichment and analysis

We used two methods to investigate the presence of circles in plants displaying the appropriate junction from the CHI target. First, to enrich any circular DNA linear DNA was depleted by incubating with Plasmid Safe Exonuclease (Epicentre) for 32 hours at 37C. The exonuclease was inactivated by incubating 30 minutes at 70C. The resulting DNA and input control DNA were used as template for PCR with GoTaq polymerase. Second, to amplify potential circular DNA sequences, 2 ul of eluted leaf genomic DNA, 16 ul of water and 1.2 ul of a single primer that is internal to the putative CH1 circles were incubated for 20 minutes at 98C, and then left to cool to room temperature. Then, 0.26 ul BSA, 2 ul of 10 mM dNTP and 2.6 ul 10X phi29 polymerase buffer were added (NEB). The reaction was split into two: A 12.5 ul reaction that received no phi29 polymerase and 11.5 ul to which 1.2 ul of phi29 polymerase (NEB) was added. The reactions were incubated for 23 hours at 30C in a thermocycler, followed by 12 hours incubation in a 30C incubator. The phi29 polymerase was heat inactivated by incubating for 10 minutes at 65C. Unidirectional amplification of a circular template with phi29 polymerase is expected to yield single stranded amplicons that may contain many copies of the circular template sequence, whereas linear templates are not expected to be amplified to more than two copies of the template. The phi29 polymerase amplicons were directly used as a template for traditional GoTaq PCR, and the resulting double stranded amplicon bands were imaged.

## Acknowledgements

This work was funded by an Innovative Genomics Institute Grant and an NSF-Plant Genome IOS Grant 144612: Rapid and Targeted Introgression of Traits via Genome Elimination. P.G.L was supported by an NSF Graduate Research Fellowship and an Elsie Taylor Stocking Fellowship.

## Author Contributions

P.G.L., S.I., K.R.A., B.R.P. and E.H.T. developed genetic constructs. M.P.A.M. designed and conducted the fluorescent microscopy experiment and localized RPS5A-tDTomato expression. P.G.L., S.I., I.M.H. and E.H.T. genotyped plants. P.G.L., S.I. and E.H.T. phenotyped plants. P.G.L. conducted the translocation and duplication experiments. P.G.L., I.M.H. and L.C. designed most of the research, analyzed data and wrote the paper.

